# TraN variants mediate conjugation species specificity of IncA/C, IncH and *Acinetobacter baumannii* plasmids

**DOI:** 10.1101/2025.11.14.688549

**Authors:** Shan He, Sophia David, Jaie Rattle, Julia Sanchez-Garrido, Josh LC Wong, Konstantinos Beis, Gad Frankel

## Abstract

IncA/C and IncH plasmids commonly carry antimicrobial resistance genes, notably *bla*_NDM-1_. Although these plasmids disseminate among Gram-negative pathogens via conjugation, the mechanisms underlying mating pair stabilisation (MPS) and conjugation species specificity remain poorly understood. In IncF plasmids, MPS is mediated by interactions between outer membrane proteins (OMP) encoded by the plasmids in the donor (TraN) and by the chromosome in the recipient. Using the Plascad database, we extracted 1,436 TraN sequences: 62.5% (898/1,436), mainly in IncF plasmids, are 550–660aa (we renamed TraN short, TraN_S_); 15% (216/1,436), in IncA/C plasmids, are 880–950aa (TraN medium, TraN_M_); and 11% (160/1,436), in IncH plasmids, are 1,050–1,070aa (TraN long, TraN_L_). A group of six TraN from *Acinetobacter baumannii* plasmids (891aa) were designated TraN V-shaped (TraN_V_). Like TraN_S_, TraN_M_ and TraN_L_ contain base and distal tip domains essential for conjugation, whereas TraN_V_ has a base and two distinct tip domains forming a V-shaped structure. TraN_M_, TraN_L_ and TraN_V_ determine conjugation species specificity, with TraN_L_ cooperating with OmpA. Tip swapping reverses conjugation specificity, revealing how TraN_M_ and TraN_L_ diversity influence plasmid host range and AMR dissemination. Our new data reveal the molecular basis of plasmid host specificity and broaden our understanding of how conjugation drives the dissemination of antimicrobial resistance genes among clinically relevant bacteria.

## Introduction

Infectious diseases pose a continuous and increasing threat to global public health (1). The rapid spread of antimicrobial resistance genes (ARGs) amongst pathogens is affecting our ability to treat many acute bacterial infections, even with last resort antibiotics (e.g. carbapenems) (2,3).

The spread of ARGs is facilitated by diverse mechanisms of horizontal gene transfer (HGT) (4,5), among which conjugation is the major driver (6). During conjugation, a plasmid is transferred unidirectionally from a donor bacterium to a recipient bacterium in a contact-dependent manner (7,8). Conjugative plasmids can disseminate within and between species, affecting bacterial evolution. Successful plasmid conjugation and maintenance in the recipient is a process involving multiple steps. First, the plasmid-encoded mating pilus bridges the donor with a potential recipient (7). In IncF plasmids, this is followed by formation of a donor-recipient synapse (8). Finally, entry of the leading single stranded plasmid DNA activates defence responses in the recipient (9), which, if counteracted by plasmid-encoded anti-defence systems, results in formation of transconjugants (10). This is followed by expression of plasmid genes encoding surface and entry exclusion (11,12).

DNA transfer is facilitated by a type IV secretion system (T4SS) (13), and the attached mating pilus. Binding of the pilus expressed by the donor to a recipient leads to mating pair formation (MPF). Based on homology between the components of the T4SS and additional accessory factors which are involved in MPF, the conjugation systems have been classified into four groups – MPFG, MPFT, MPFI and MPFF (14,15). MPFF plasmids, which express long conjugative pili and belong to the IncF, IncH and IncA/C groups, account for more than a third of conjugative plasmids isolated from Gammaproteobacteria and are particularly highly represented amongst Enterobacterales species (16).

The reference IncHI1 plasmid, R27 was isolated from *Salmonella enterica* serovar Typhi in 1961 (17). IncHI1 and IncHI2 plasmids are thermosensitive for transfer with an optimum rate of transfer between 22 and 30°C (18). A mutation in *htdA* derepresses R27 (drR27) resulting in constitutive expression of the conjugation machinery and the H pilus (19). The reference IncA/C plasmid, RA1, was isolated from *Aeromonas liquefaciens* in 1971 (20). RA1 expresses the conjugation machinery constitutively at 30°C (21).

The cryo-EM structures of the mating pili expressed by the IncF-like plasmids pOX38, pEB208 and pKpQIL revealed that the linear TraA pilin subunits form helical assemblies with phosphatidylglycerol (PG) molecules at a stochiometric ratio of 1:1 (22). More recently we reported that the H pilus, encoded by R27, is an assembly of PG-associated cyclic ThrA pilin subunits (23). Pilus-mediated MPF enables inefficient DNA transfer from a distance (24). Formation of MPF is followed by pilus retraction, leading the donor and recipient bacteria forming mating junctions (25), characterised by intimate wall-to-wall contact, through a process termed mating pair stabilisation (MPS) (26), which mediates efficient DNA transfer (24). We have reported that in IncF-like plasmids, MPS is mediated by interactions between the plasmid-encoded one of four TraN isotypes in the donor, classified as TraNa, TraNb, TraNg and TraNd, specifically with OmpW, OmpK36, OmpA, OmpF in the recipient, respectively (24).

AlphaFold structural prediction of the different TraN isotypes encoded by IncF-like plasmids reveal that they consist of a conserved base domain, predicted to embed the protein in the donor outer membrane and a variable distal tip domain which mediates conjugation efficiency, specificity and host range (27). We have shown that while TraNα and TraNδ mediate broad conjugation host range, TraNβ and TraNγ mediate narrow conjugation host range as they are mainly found in *Klebsiella* species and in *E. coli* respectively (24).

While IncA/C and IncH plasmids also belong to the MPFF group and encode TraN, less is known about how they establish donor-recipient interactions and MPS, even though they are important vehicles for the spread of antibiotic resistance in *Escherichia coli, Klebsiella pneumoniae* and *Acinetobacter baumannii*, particularly the carbapenemase New Delhi metallo-b-lactamase-1 (NDM-1) (28).

In this study we characterised the IncA/C and IncH plasmid-encoded TraN, as well as TraN encoded by plasmids found exclusively in *A. baumannii*. As these TraN proteins are ca.50% larger than IncF plasmids-encoded TraN, which we have now reclassified as TraN short (TraN_S_), we classified the IncA/C-encoded TraN as TraN medium (TraN_M_), the IncH-encoded TraN as TraN long (TraN_L_) and the *A. baumannii-*encoded TraN as TraN V-shaped (TraN_V_), respectively. Using bioinformatic, structural, and conjugation analyses, we reveal their diversity, host range preferences, and receptor dependencies. Together, these findings provide new insights into the molecular basis of plasmid host specificity and broaden our understanding of how conjugation drives the dissemination of antimicrobial resistance among clinically relevant bacteria.

## Results

### Identification of TraN derivatives in IncA/C and IncH plasmids

We searched for TraN protein sequences among 1517 MPFF plasmids in the Plascad database (29), using the sequence annotations. 89.5% (1358/1517) of the plasmids contained a single TraN and 2.5% (38/1517) contained 2-3 copies. No TraN was found in 8.0% (121/1517) of the plasmids. Altogether we extracted 1436 TraN protein sequences from the 1517 plasmids.

The length of the identified TraN proteins was highly variable, 62.5% (898/1436) were 550-660 amino acids (aa) long, which we have now renamed TraN short (TraN_S_), with many of these derived from plasmids carrying IncF replicons and carried by Enterobacterales species. We also noted additional distinct groupings based on size, with a further 15.0% (216/1436) possessing a length of 880-950 aa, which we classified TraN medium (TraN_M_), and 11.1% (160/1436) constituting 1050-1070 aa, which we classified TraN long (TraN_L_). Other smaller groups include six TraN of 891 aa from *A. baumannii* plasmids, which we classified as TraN V-shaped (TraN_v_), nine TraN of 1882 aa from different *Xanthomonas* species, and nine TraN of 885-931aa from different Enterobacterales species.

We compared the pairwise protein similarity of the 400 TraN proteins ≥880 aa (Fig. 2). A heatmap showing these pairwise similarities illustrates the high level of diversity found among these proteins overall, with similarities as low as 10% between some pairs. However, two major groups, which can be further subdivided into subgroups, were observed based on protein similarity (**Fig. 1)**. The first major group comprises TraN_M_, which is largely associated with either an IncA/C (2/184) or IncA/C2 replicon (151/184), including the prototypical IncA/C plasmid RA1. The other major group, TraN_L_, is largely associated with different IncH replicons (135/155), including the prototypical IncHI1A plasmid R27. The other smaller groups represented in the heatmap include TraN_V_ (**Fig. 1**).

**Figure 1.**
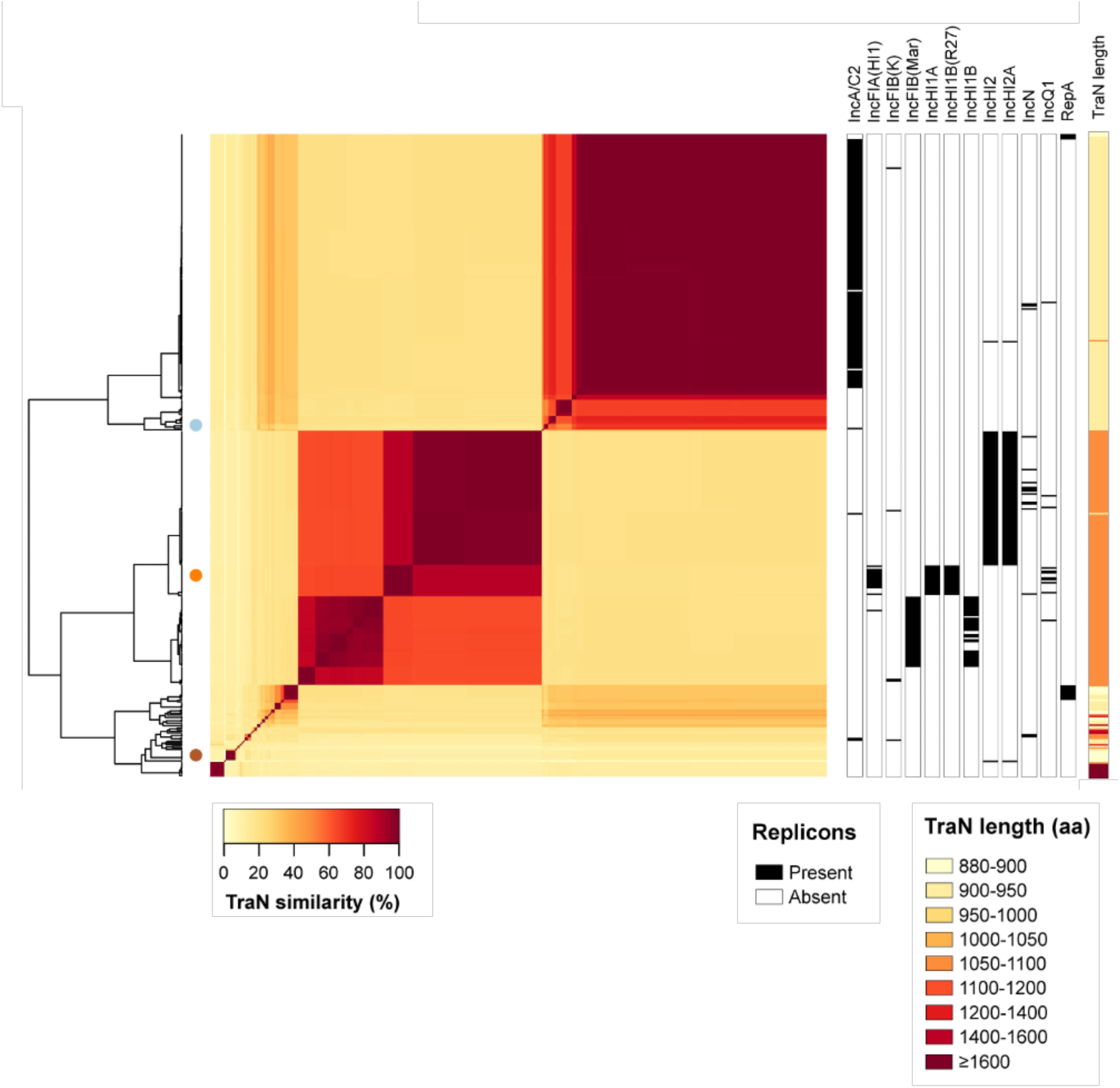
Pairwise similarity of TraN proteins. Heatmap showing the pairwise similarity among 392 “long” TraN protein sequences (≥880aa). Metadata columns show the TraN protein length (aa) and the presence/absence of particular plasmid replicons (those found in ≥5 plasmids) among the associated plasmids.

**Figure 2.**
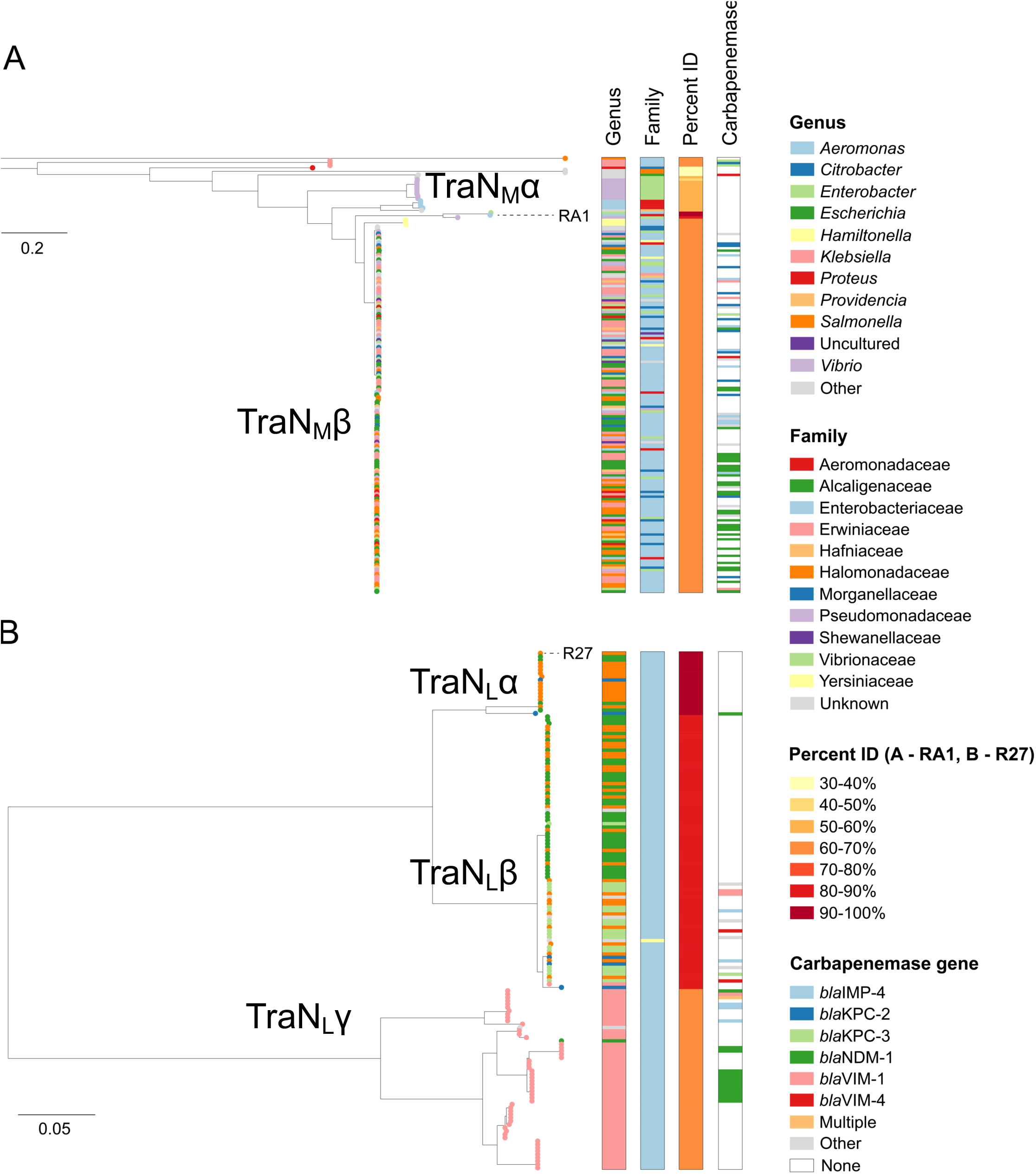
Phylogenetic analysis of TraN proteins. Phylogenetic trees of “long” TraN protein sequences (≥880aa) with ≥30% similarity to TraN(RA1) (**A**) and TraN(R27) (**B**). The trees contain 184 and 155 sequences, respectively, and are midpoint-rooted. The tree tips show the host genus of the associated plasmid. Metadata columns show the host genus and family of the associated plasmid, the percent identity of each TraN variant with respect to TraN(RA1) (A) or TraN(R27) (B), the presence of IncA/C replicons (A) or IncH replicons (B) and the presence of carbapenemase gene variants within the plasmid. The scale bars show the number of substitutions per site. Similar visualisations can be accessed via Microreact: https://microreact.org/project/tran-ra1 (A) and https://microreact.org/project/tran-r27 (B).

### Distribution of TraN_M_ and TraN_L_ in host taxa

To investigate TraN_M_ and TraN_L_ relatedness at high resolution and their association with host taxa, we constructed maximum-likelihood phylogenies of all TraN proteins with ≥30% similarity to RA1-and R27-encoded TraN. This revealed the TraN_M_ group can be divided into a minor TraN_M_α, which includes RA1, and a major TraN_M_β clades. The TraN_M_ proteins represented in the tree were obtained from plasmids derived from at least 11 different host families, with Enterobacterales accounting for 66.3% (122/184). TraN_M_ variants from different families (and different genera) were highly interspersed within the tree, suggestive of frequent exchange of the associated plasmids between these host taxa (Fig. 2A; https://microreact.org/project/4wtmDXd6nPqktyE8frk3J2-tran-30percent-aa-similarity-to-ra1).

The TraN_L_ group can be divided into three clades we named TraN_L_α, which includes R27, TraN_L_β and TraN_L_γ. Notably, all but one of the TraN_L_ proteins are from plasmids found in Enterobacterales. Of note, we observed clear associations of clades with one or more genera. Plasmids encoding TraN_L_α are mainly found in *Salmonella*/*Escherichia*, TraN_L_β are mainly found in *Salmonella*/*Enterobacter* and plasmid encoding TraN_L_γ are almost exclusively found in *Klebsiella* plasmids, (Fig. 2B; https://microreact.org/project/vkQNTtANTwjv9zNgpqybwe-tran-30percent-aa-similarity-to-r27).

Importantly, 39.1% (72/184) and 15.5% (24/155) of the plasmids encoding TraN_M_ and TraN_L_ carry a carbapenemase gene, respectively, with a particular predominance of *bla*_NDM-1_ (Fig. 2). Taken together, these findings suggest that a large proportion of plasmids expressing TraN_M_ encode *bla*_NDM-1_ and exhibit a broad host range while a small proportion of plasmids expressing TraN_L_ encode *bla*_NDM-1_ and show high degree of host restriction.

### TraN_M_ and TraN_L_-mediated recipient conjugation specificity

We have shown before that the tips of the four TraN_S_ variants (TraN_S_ α, β, γ, δ), (Fig. 3A), mediate MPS, conjugation efficiency and species specificity (24). The AlphaFold structural predictions of RA1 TraN_M_ and R27 TraN_L_ revealed that their structures can also be divided into base and tip domains, with the most prominent differences in the size and conformation of their tips (Figure 3B-C).

**Figure 3.**
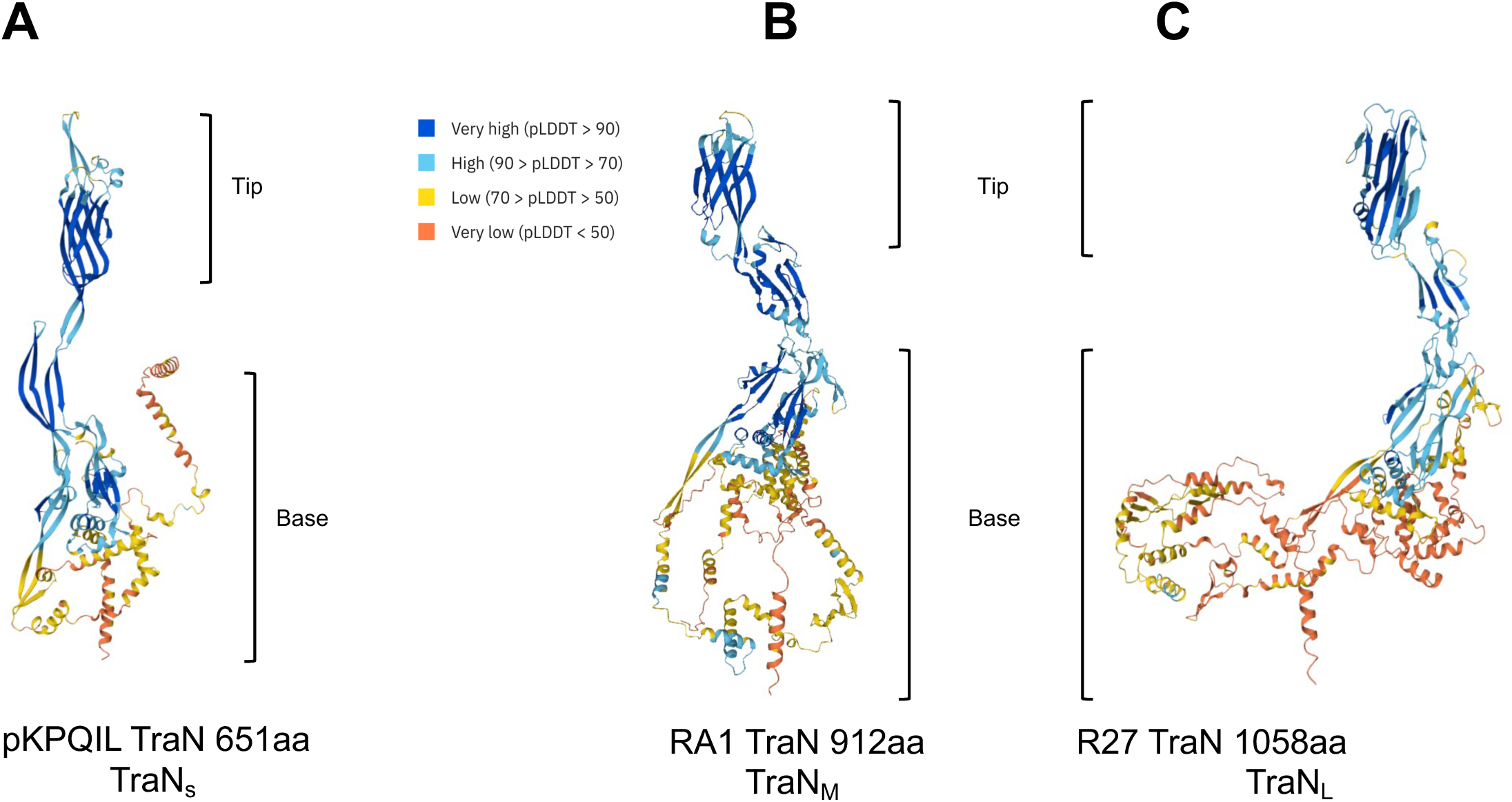
Alpha-fold prediction and conjugation specificity of R27 and RA1. AlphaFold structural prediction of TraN encoded by pKpQIL (A), RA1 (B) and R27 (C). The predicted local distance difference test (pLDDT) is a per-residue measure of local confidence, scaled from 0 to 100.

We used *E. coli* MG1655 containing either R27 or RA1 as donors and six well-characterized pathogenic species from the *Enterobacterales* family as recipients in conjugation assays, MG1655 was also used as a recipient control. In agreement with the observed distribution of TraN_M_ in clinical isolates (Fig. 2), RA1 was conjugated efficiently into *Citrobacter freundii (C. freundii), Shigella sonnei (S. sonnei)*, and MG1655. Conversely, RA1 was conjugated with lower efficiency (∼80-fold lower) into *K. pneumoniae*, *Enterobacter cloacae (E. cloacae)*, and *Salmonella enterica* serovar Enteritidis (*S. enteritidis*) and with a very low frequency into enteropathogenic *E. coli* (EPEC, strain E2348/69) (Fig. 4A). R27 plasmid was conjugated with a high frequency into *K. pneumoniae*, EPEC and MG1655 and with lower frequency into *E. cloacae*, and *S. sonnei* (Fig. 4B). The conjugation frequency dropped significantly when *S.* Enteritidis and *C. freundii* were used as recipients. We have previously shown that the tip of TraN_S_ provides species specificity. Based on this we sought to establish if the tips of R27 and RA1 also mediate distinct conjugation species specificity among Enterobacterales. To this end we first generated RA1 and R27 tip-less mutants (Δtip) to be used as negative controls. Quantifying conjugation frequency into MG1655 revealed that RA1 and R27 Δtip were conjugated at ca. 350-fold lower efficiency compared to the corresponding wild-type (WT) plasmids (Fig. 4C-F). We then swapped the tips between RA1 and R27 and quantified conjugation frequencies into *K. pneumoniae* and *S. sonnei* recipients as representatives. This revealed that compared to R27, the R27::TraN_RA1_ was now conjugated with lower efficiency into *K. pneumoniae* and higher efficiency into *S. sonnei.* Conversely, compared to RA1, the RA1::TraN_R27_ conjugated with higher efficiency into *K. pneumoniae* and lower efficiency into *S. sonnei.* (Fig. 4F).

**Figure 4.**
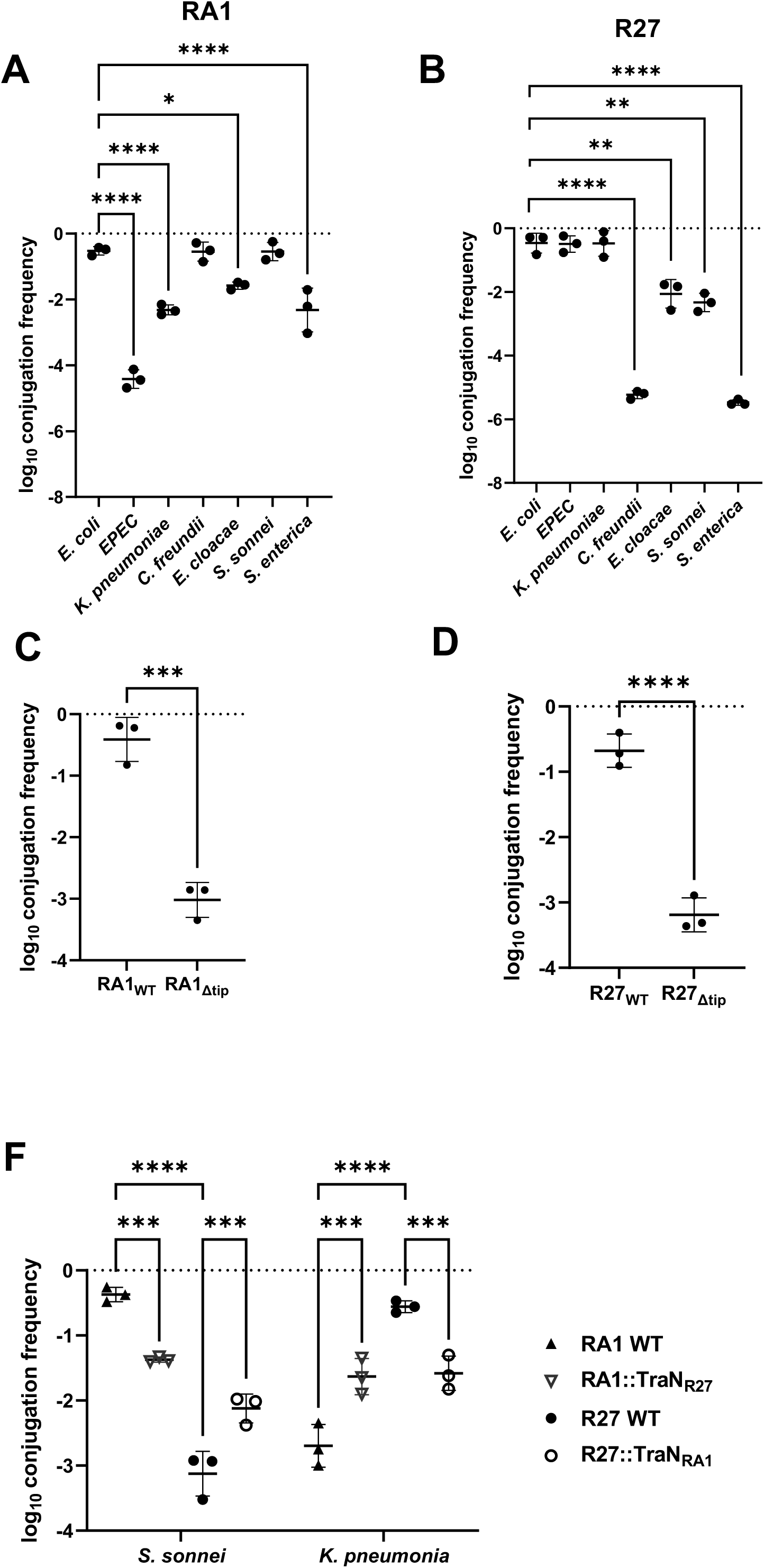
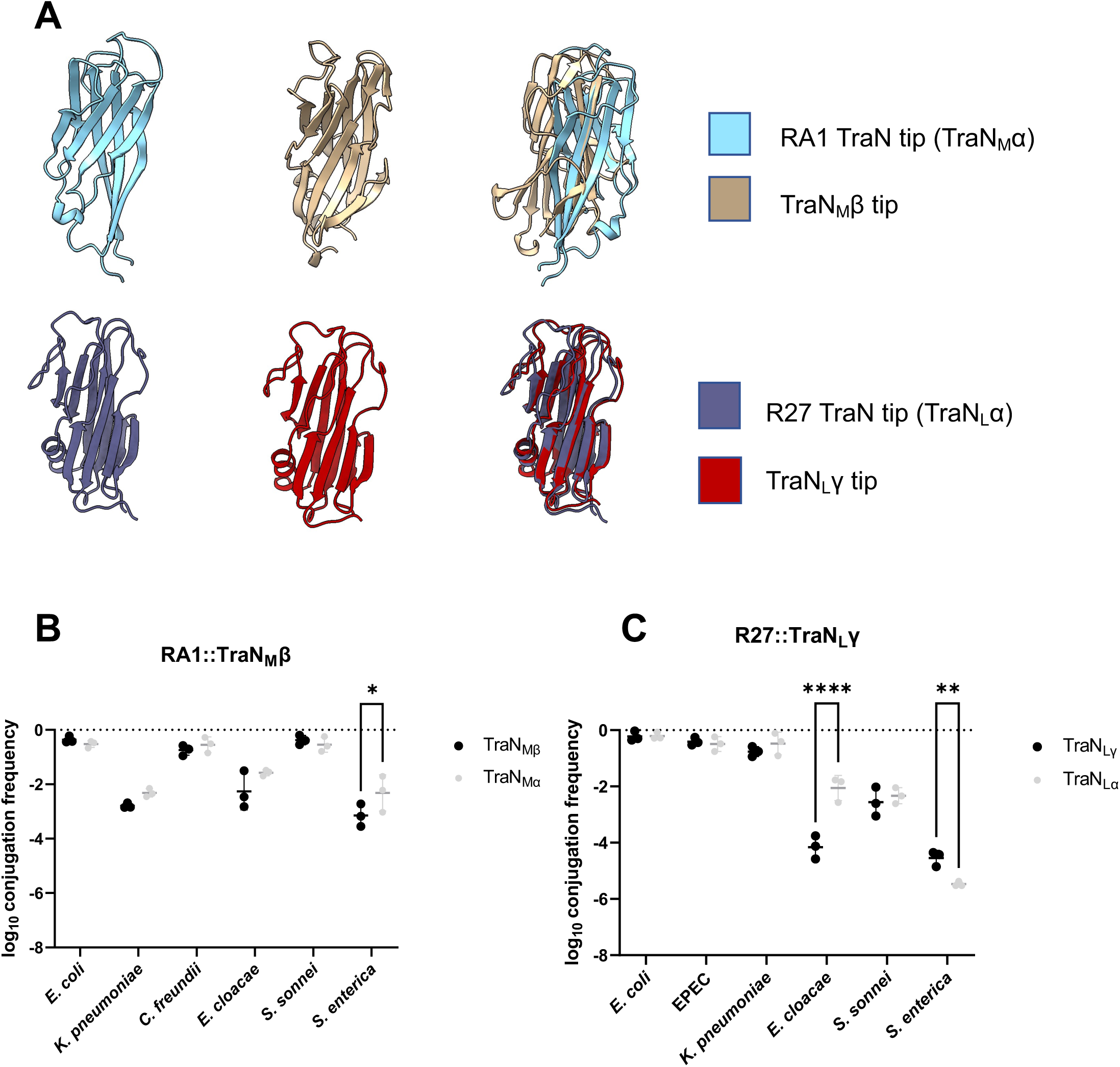
TraN_M_ and TraN_L_-mediated recipient conjugation specificity. Conjugation of RA1 (A) and R27 (B) into different *Enterobacterales and E. coli* MG1655. Log conjugation frequency data are presented as mean□±□s.d. of three biological repeats, analysed by repeated measures one-way ANOVA with Tukey’s multiple comparison test. (C) schematic representation of the RA1 and R27 TraN domains. Tip-less RA1 (D) and R27 Log conjugation frequency data are presented as mean□±□s.d. of three biological repeats, analysed by a two-sided paired t-test. (E) TraN swaps between RA1 and R27 reverses conjugation specificity into *S. sonnei* and *K. pneumoniae* recipients. All the data has been log transformed, and presented as mean□±□s.d. of three biological repeats, analysed by repeated measures two-way ANOVA with Tukey’s multiple comparison test. *, *P<0.05*; **, *P<0.01*; ****, *P<0.0001*.

We also tested if the TraN_M_ and TraN_L_ variants seen in the phylogenetic tree (Fig. 2) affected conjugation species specificity. To this end we selected two resistance plasmids, pNDM-US (30), encoding TraN_M_β and pNDM-MAR (31), encoding TraN_L_γ (Fig. 2). AlphaFold structural prediction showed that the structure of the TraN_L_ tips encoded by R27 and pNDM-MAR are very similar despite variations in their sequences (Fig. 5A). In contrast, the TraN_M_ tips encoded by RA1 and pNDM-US showed different folds, particularly at the distal loops (Fig. 5A).

**Figure 5.**
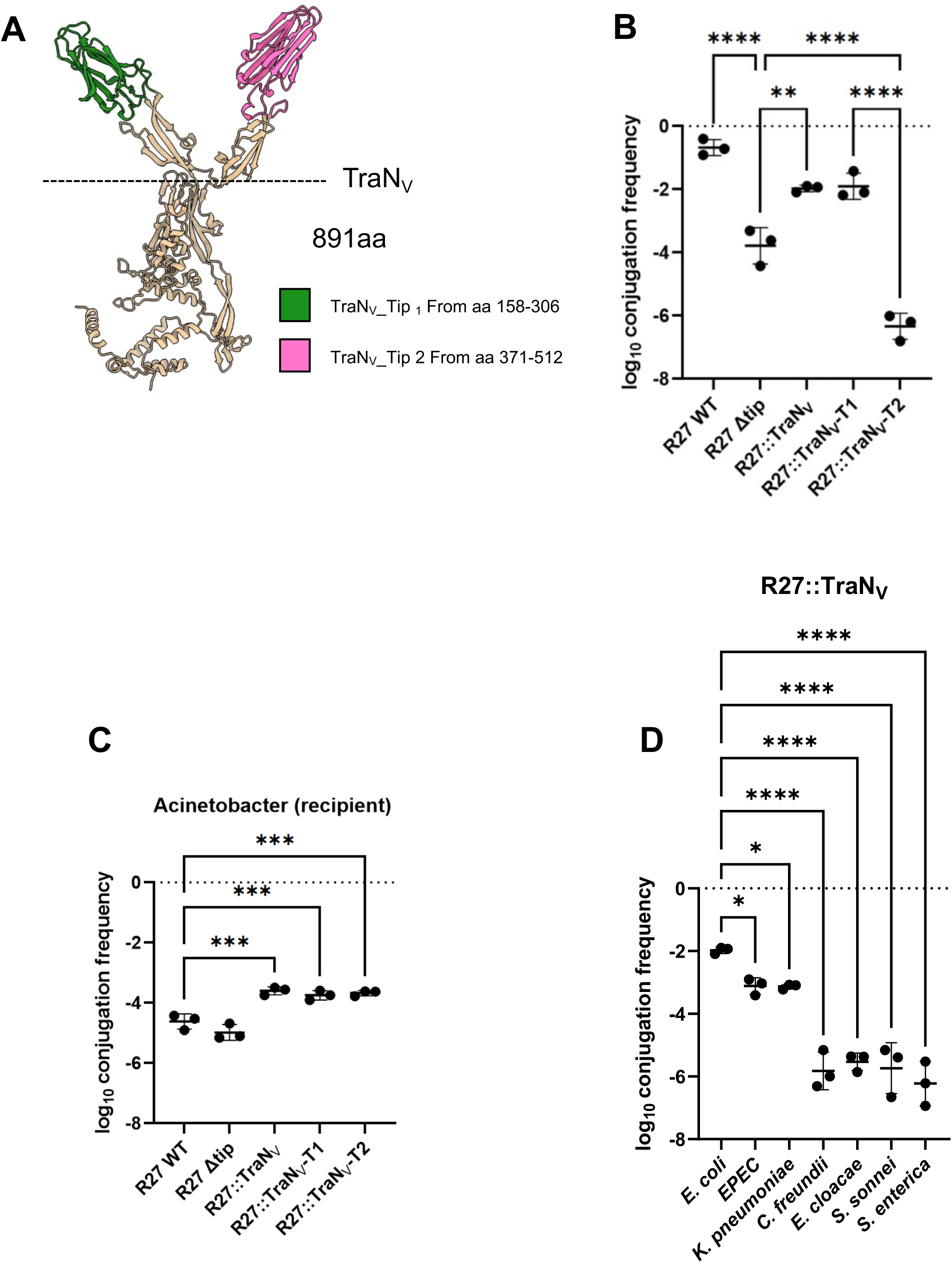
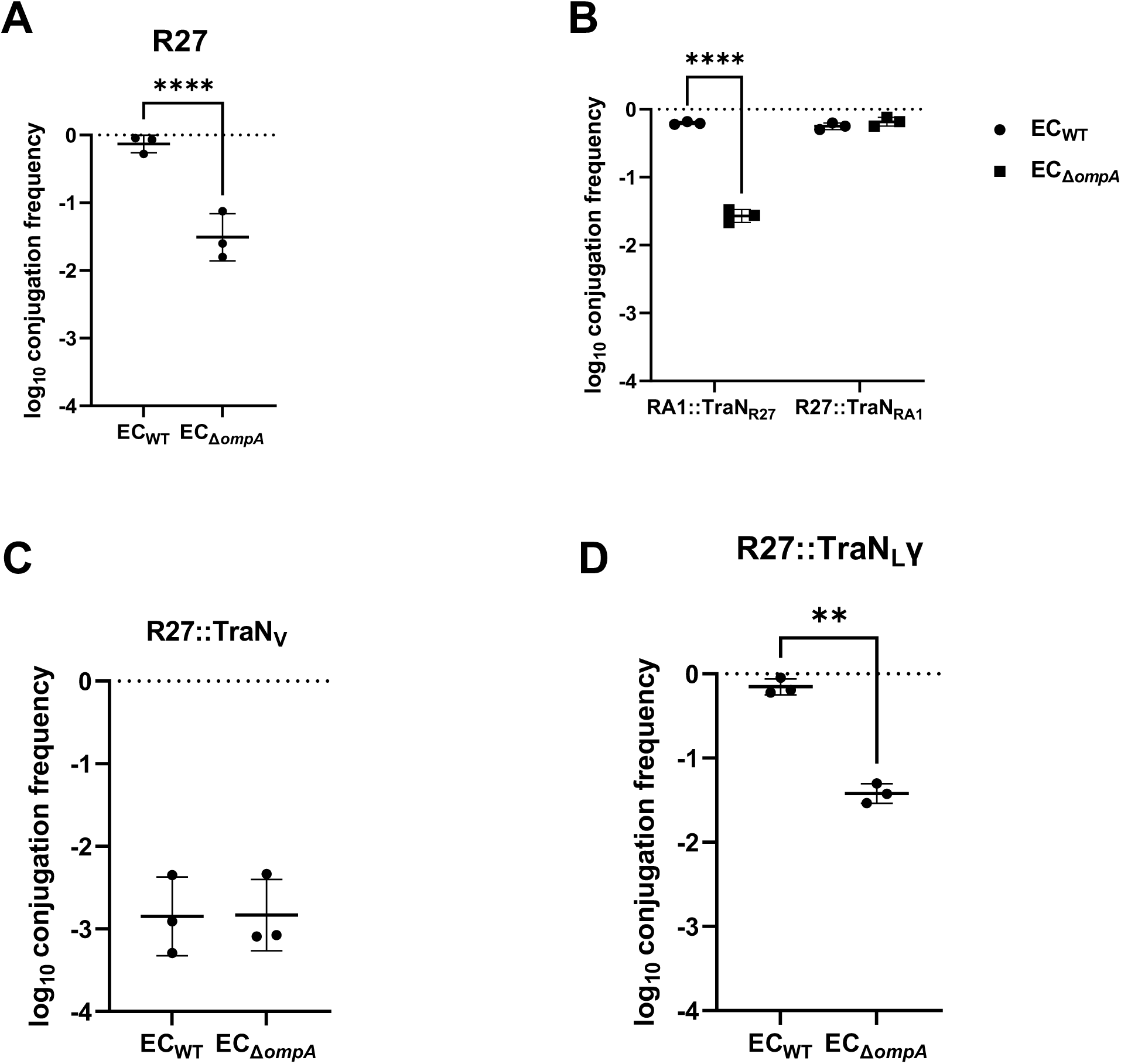
Conjugation specificity of TraN_M_β and TraN_L_γ. (A). Cartoon representation of the AlphaFold structural prediction of the TraN variant tips. The TraN_M_ and TraN_L_ subfamilies display structural similarities between them. Comparison of conjuation species specificity between TraN_M_α and TraN_M_β (D) and TraN_L_α and TraN_L_γ (E). The comparison was performed between the TraN variant and their prototypical TraN, and the data has been log transformed, and presented as mean□±□s.d. of three biological repeats, analysed by repeated measures two-way ANOVA with Tukey’s multiple comparison test. ns, non-significant; *, *P<0.05*; **, *P<0.01*; ***, *P<0.001*, ****, *P<0.0001*.

Our panel of Enterobacterales species as recipients revealed that despite the structural differences, the TraN_M_β tip mediated similar conjugation efficiency as the TraN_M_α tip except a subtle decrease of conjugation into the *S.* Enteritidis recipient (Fig. 5B). In contrast, while structurally conserved, R27 expressing the TraN_L_γ tip mediated lower conjugation efficiency specifically into the *E. cloacae* recipient and higher conjugation frequency into the *S.* Enteritidis recipient compared with TraN_L_α; conjugation efficiency into other recipients did not change (Fig. 5C).

Taken together these results show that the TraN_M_ and TraN_L_ tips mediate MPS and recipient conjugation specificity like the TraN_S_ (24) Despite the high degree of structural similarity between the TraN_L_ variants, subtle sequence changes affect conjugation efficiency into specific recipients.

### TraN_V_ mediates conjugation into *A. baumannii* and selected Enterobacterales species

We identified six TraN-encoding plasmids in *A. baumannii* (Fig. 1), whose AlphaFold structural prediction revealed a unique architecture of a base domain presenting two structurally distinct tips, arranged into a V-shape; we named them TraN_V_ and annotated the respective tips as T1 and T2 (Fig. 6A). To investigate the role TraN_V_ plays in conjugation, we ‘transplanted’ the V-shaped tip and individually T1 and T2 from pABAY10001, an *A. baumannii* 54 kb plasmid containing no antibiotic resistance genes (GenBank: MK386682.1), onto the R27 TraN base.

**Figure 6.** Characterisation of TraN_V_. AlphaFold structural prediction of TraN_V_. TraN_V__Tip 1 and TraN_V__Tip 2 are shown in green and pink colours respectively. The dash line marks for the V-shape 2 tips region, that was transplanted onto R27 (A). Conjugation of TraN_V_ and its respective tips into *E. coli* MG1655 recipient (B). Conjugation of TraN_V_ and its respective tips from E. coli donor to *A. baumannii* recipient, using WT and Δtip R27 as controls (C). Conjugation species specificity of TraN_V_, transplanted onto the R27 TraN_L_ base (D). All the data has been log transformed and presented as mean□±□s.d. of three biological repeats, analysed by repeated-measures One-way ANOVA with Tukey’s multiple comparison test using R27 WT as a control. *, *P<0.05*; **, *P<0.01*; ***, *P<0.001*, ****, *P<0.0001*.

We first used MG1655 containing these chimeric TraN variants as donors and MG1655 and an *A. baumannii* strain (ATCC17978) as recipients. MG1655 donors containing R27 or R27Δtip were used as controls (Fig. 6B). R27::TraN_V_ and R27::TraN_V_-T1 were conjugated with similar efficiencies into MG1655, intermediate between R27 and R27Δtip. In contrast, R27::TraN_V_-T2 was not conjugated (Fig. 6B). In contrast, R27::TraN_V_, R27::TraN_V_-T1 and R27::TraN_V_-T2 were conjugated with similar efficiencies into *A. baumannii*, which were higher than both R27 and R27Δtip (Fig. 6C). This suggests that that while TraN_L_-does not mediated conjugation into *A. baumannii*, TraN_V_-T1 is the dominant tip during TraN_V_-mediated conjugation into *E. coli*. In contrast, TraN_V_, TraN_V_-T1 and TraN_V_-T2 are equally functional in conjugation into an *A. baumannii* recipient.

Finally, we investigated the conjugation host range mediated by R27::TraN_V_ using our Enterobacterales panel as recipients. This revealed that R27:TraN_V_ was conjugated in high efficiency into MG1655, EPEC and *K. pneumoniae* and much lower efficiency into *C. freundii*, *E. cloacae*, *S. sonnei*, and *S.* Enterica (Fig. 6D).

### TraN_L_-mediated conjugation is dependent on OmpA

We hypothesised that like TraN_S_, MPS is mediated by cooperation between the tip domains of TraN_M_, and TraN_L_ in the donor and an OMP in the recipient. To test this hypothesis, we used a selection of *E. coli* OMP deletion mutants as recipients (Fig. S1). This has shown that none of the tested mutations affected conjugation efficiency of RA1 (Fig. S1A). In contrast, R27 was conjugated at lower frequency specifically into an Δ*ompA* recipient (Fig. 7A, Fig S1B). To validate this, we used the swapped RA1 and R27 tips and the Δ*ompA* recipient, and we also included R27::TraN_V_. This revealed that conjugation of R27-TraN_M_ and R27::TraN_V_ was OmpA-independent, while conjugation of RA1-TraN_L_ was now OmpA-dependent (Fig. 7B-D). We concluded that TraN_L_α-mediated MPS is dependent on OmpA while the partner OMP of TraN_M_ and TraN_V_ remain unknown.

**Figure 7.** Conjugation of TraN_L_ is OmpA-dependent. Conjugation of R27 is OmpA-dependent (A). Swapping tips between RA1 and R27 reverses OmpA dependency (B). OmpA-dependent experiment for TraN_V_ (C) TraN_L_γ in R27maintained the OmpA-dependency (D).

## Discussion

The spread of ARGs is at the heart of the silent AMR pandemic (32). IncA/C and IncH plasmids frequently carry carbapenemsases; indeed, a recent study described the spread of an IncH imipenemase encoding plasmid among diverse Enterobacterales species between 2016 and 2019 across a London regional network. (33) Yet, little is known about MPS and the mechanism of transfer of IncA/C and IncH plasmids.

Bioinformatics analysis shows that all the TraN proteins encoded by IncA/C plasmids fall into the TraN_M_ category. While we identified the OMP partners for the TraN_S_ and TraN_L_ variants, the partner of TraN_M_ remains unknown. The TraN_M_ can be divided into two clades, a small TraN_M_α cluster which includes the prototype plasmid RA1, and a larger TraN_M_β clade, which includes the pNDM-US plasmid, isolated in the USA from *K. pneumoniae* and carries the *bla_NDM_* gene (30). Considering the bias in genome sequencing towards strains recovered from sever clinical cases, this suggests that TraN_M_β is more commonly found in these isolates. While most plasmids in this cluster were found in Enterobacterales species, they were also associated with other families, suggesting that IncA/C are broad host range plasmids. Our conjugation assays revealed that IncA/C plasmids are conjugated efficiently into the representative Enterobacterales species we used, except into EPEC, possibly due to expression of defence systems (34).

All the TraN proteins expressed by IncH plasmids belong to the TraN_L_ category. AlphaFold predictions indicate that despite sequence and host range variabilities, the TraN_L_ variants share similar structure and use OmpA as a recipient partner despite sequence differences. In contrast to TraN_M_, with only one exception, all TraN_L_ proteins have been isolated from Enterobacterales species, suggesting that IncH plasmids mainly circulate specifically within this family. Analysis of the IncHI1B pNDM-MAR plasmid, isolated in the USA from *K. pneumonia* and carrying the *bla_NDM_* gene (31), revealed that its TraN belongs to the TraN_L_γ variant. While the TraN_L_γ is found almost exclusively in *K. pneumoniae*, it can mediate efficient DNA transfer into other Enterobacterales species, including EPEC, raising the possibility that additional factors on plasmids endogenously encoding TraN_Lγ_ compatibility of defence and anti-defence systems (34), allow their replication or maintenance specifically in *Klebsiella*.

In addition to the TraN_S_, TraN_M_ and TraN_L_, we also found a small TraN_V_ clade, unexpectedly encoded by plasmids found in *A. baumannii* in China and South Korea. Structural predictions revealed that the TraN_V_ base domain is extended by two distinct tips. While a chimera of TraN_V_, T1 or T2 on the R27 base resulted in similar conjugation efficiency into *A. baumannii* recipient, only TraN_V_ and T1 were functional when *E. coli* was used as a recipient. This suggests that while the receptor for T1 is shared between the two recipients, the receptor for T2 is only present in *A. baumannii.* Testing conjugation of TraN_V_ into multiple species revealed highest efficiency into *E. coli, K. pneumoniae* and EPEC, suggesting that these species could acquire such plasmids. Conjugation into the other species was similar to the level recorded for the Δtip, suggesting that these species have a relatively low likelihood of acquiring TraN_V_-encoding plasmids in the future.

In summary, this study focused on TraN_M_, TraN_L_, and TraN_V_, deepening our understanding of the mechanism of spread of IncA/C, IncH and the TraN_V_-encoding plasmids in Enterobacterales species. Combining our data of TraN-mediated species specificity could potentially be used as predictors for future plasmid transmission.

## Acknowledgements

We thank Prof. Ruth Hall, University of Sydney and Dr. Choong-Min Ryu, Korea Research Institute of Bioscience and BioTechnology for gifting us RA1 and pABAY10001, respectively. We thank Prof. John Tregoning for the *Acinetobacter baumannii* strain. GF is supported by a Wellcome Trust Investigator Award grants 224282/Z/21/Z.

## Author contributions

GF and KB conceived and supervised the study. SD performed the bioinformatics. JR, JS-G and JLCW provided technical support.

## Methods

### Identification of TraN homologues and characterisation of the associated plasmids

We used the Plascad database (29) to identify publicly-available plasmid sequences possessing the mating pair formation “F” (MPFF) system. We then identified and extracted protein sequences from the associated genebank files that were annotated as: product = “traN” OR “TraN” OR “trhN” OR “TrhN”, or gene = ““traN” OR “TraN” OR “trhN” OR “TrhN”. We used Clustal Omega v1.2.4 (35) to align the resulting protein sequences and generate a pairwise similarity matrix. We generated a heatmap showing the pairwise similarities between all “long” TraN proteins (≥880aa) using the heatmap.2 function from the gplots v.3.2.0 package in RStudio (R v4.4.2).

Plasmid replicon types were identified from the MPFF plasmids using PlasmidFinder (database version 29/11/2021) (36). Carbapenemase genes were identified using AMRFinderPlus v3.10.23, database version 2021-12-21.114 (37).

### Phylogenetic analysis of TraN homologues

Separate alignments were generated incorporating all TraN protein sequences of ≥880aa with ≥30% similarity to those from the RA1 and R27 plasmids, respectively. ModelTest-NG v0.2.0 (38) was used to select the most suitable evolutionary model to use in each phylogenetic analysis. RAxML-NG v1.2.0 (39) was used to generate a maximum likelihood phylogenetic tree with 1000 bootstrap replicates from each alignment. The phylogenetic trees were visualised together with metadata using Microreact (40).

### Bacterial strains and plasmids

The bacterial strains, conjugative plasmids and mutagenesis vectors used are listed in Supplementary Table 1, 2 and 3 respectively. Unless otherwise stated, bacteria were cultured in Lysogeny Broth (LB) at 37□°C, 200 r.p.m. When needed, antibiotics were used at the following concentrations: chloramphenicol (30□μg/ml), streptomycin (50□μg/ml), kanamycin (50μg/ml), gentamicin (10□μg/ml), tetracycline (10μg/ml).

### Conjugation assay

All the conjugative plasmids are expressed in trp-*E. coli* MG1655, our conjugative “donor”. Overnight cultures of donor and recipient strains were washed in fresh LB and mixed in a ratio of 10:1 (R27) or 2:1 (RA1). This mixture was further diluted 1 in 25 in fresh LB and 40 μl spotted onto LB agar plates. Plates were subcultured for 2-3 hours at 37°C before incubation at 25□ (R27) or at 30 ℃ overnight (RA1). The conjugation mixture was collected and resuspended in 1 ml of sterile PBS. The transconjugants were selected by plating the conjugation mixture onto a M9 minimal media agar plate containing the appropriate selection marker (chloramphenicol for R27; tetracycline for RA1). Conjugation frequency was calculated as the ratio of the colony forming units per ml (CFU/mL) of transconjugants to the CFU/mL of recipients, and the data were log_10_ transformed before statistical analysis.

### Generation of mutants

All genomic mutations were made in *E. coli* MG1655, using a two-step recombination methodology. Mutagenesis vectors were mobilized from *E. coli* CC118λpir into pACBSR-carrying strains through a tri-parental conjugation using the E. coli 1047 pRK2013 helper strain. Merodiploid colonies were selected on LB agar containing gentamicin and streptomycin. Selected colonies were grown for at least 4□h in LB supplemented with streptomycin and 0.4% l-arabinose to induce expression of the I-SceI endonuclease from the pACBSR plasmid. Cultures were streaked onto LB agar containing streptomycin and screened for the intended mutations. Mutations in plasmids were introduced using the same methodology.

Mutagenesis vectors were generated by Gibson Assembly (New England Biolabs, E2611L) using the pSEVA612S backbone and were maintained in *E. coli* CC118λpir cells. Site-directed mutagenesis on previously generated vectors was performed according to the Q5 Site-Directed Mutagenesis Kit protocol (New England Biolabs, M0554S). Primers used to generate the mutagenesis vectors and for screening are listed in Supplementary Table 4. All mutations were confirmed by sequencing (Eurofins).

### Generation of TraN AlphaFold models

In the absence of homologous TraN structures, *ab initio* models were generated by AlphaFold (41). TraN sequences were submitted to the AlphaFold Colab server with the default settings; the signal peptide was removed from all sequences before modelling. Each structural model was validated by analysing the confidence score as generated by the pLDDT. Molecular graphics and superimposition analysis were performed in UCSF ChimeraΧ-1.8 (42).

### Statistics and reproducibility

All data are representative of 2-4biological independent repeats. Statistical analyses were performed on GraphPrism 9.5.0 (GraphPad software) and data was checked for normality after Log10 transformation using the Shapiro-Wilk test. Conjugation data were analysed by repeated measures, one-way ANOVA with Tukey’s or Dunnett’s multiple comparison test, as appropriate (detailed in the figure legends). Where only two recipient strains were being compared, a two-sided paired t-test was used. P values less than 0.05 were considered significant.

### TraN variants sequences

TraN sequences were obtained from the following reference plasmids: R27 (accession ID: AF250878), RA1 (accession ID: FJ705807), p-NDM-US (accession ID: CP006661), p-NDM-MAR (accession ID: JN420336), pABAY10001 (accession ID: MK386682).

**Table 1.** List of TraN proteins

| **TraN** | **Plasmid** | **Protein ID** | **Tip residues** |
| --- | --- | --- | --- |
| TraN<sub>M</sub>α | RA1 | <u>ACN66968.1</u> | 230C-360D |
| TraN<sub>M</sub>β | pNDM-US | <u>AHI38890.1</u> | 205N-354V |
| TraN<sub>L</sub>α | R27 | <u>AAF69844.1</u> | 305S-457E |
| TraN<sub>L</sub>β | pHNAH67 | <u>APZ78090.1</u> | 305T-457E |
| TraN<sub>L</sub>γ | pNDM-MAR | <u>AFB82831.1</u> | 304S-457D |
| TraN<sub>V</sub> | ABAY10001 | <u>QBN23304.1</u> | 149V-628R |

