## Supplementary material for "TraN variants mediate conjugation species specificity of IncA/C, IncH and *Acinetobacter baumannii* plasmids"

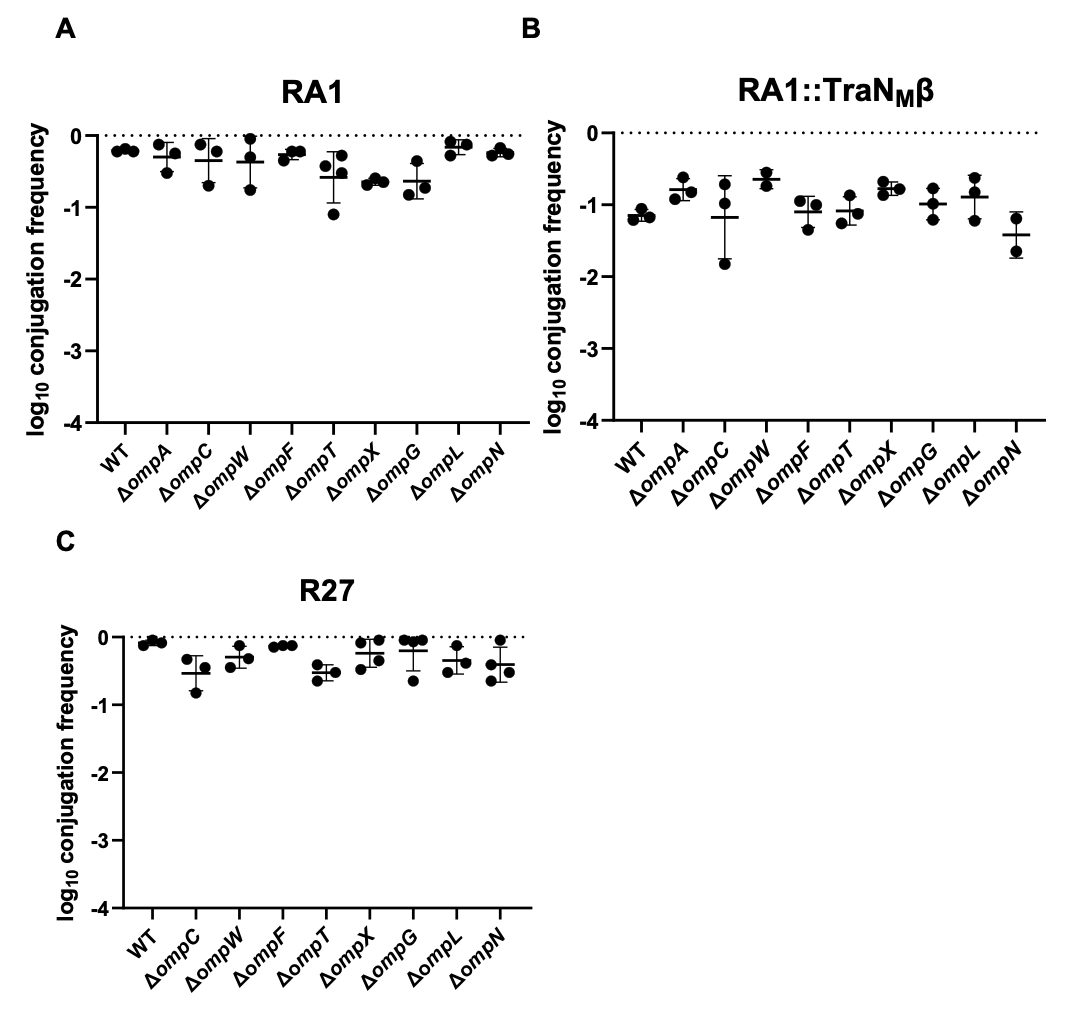

**Supplementary Figure 1.** Outer-membrane protein mutants from KEIO collection to test the potential receptors for TraN_M_α, TraN_M_β, and TraN_L_α respectively. All the data has been log transformed and presented as mean ± s.d. of at least two biological repeats, analysed by repeated-measures One-way ANOVA with Tukey’s multiple comparison test using *E. coli* MG1655 WT as a control. No significant differences were found among any mutant.

**Supplementary Table 1. Bacterial strains used in this study**

| Strain | Description |
| --- | --- |
| CC118 λ pir | Expresses the Pi protein for the replication of plasmids with the R6K origin. |
| *E. coli* 1047 pRK2013 | Triparental conjugation helper strain. Kanamycin resistant |
| *E. coli* MG1655 *trp*- | Donor strain for conjugation assays. It is *trp*- mutant, won’t be able to grow on the minimal media. |
| *E. coli* MG1655 | *E. coli* K-12 strain |
| EPEC | *Enteropathogenic E. coli* e2348/69 |
| *K. pneumoniae* | ICC8001. Parental wild type (WT) strain of K. pneumoniae ATCC43816 serially passaged in vitro on Rifampicin(100μg/ml) followed by two passages in BALB/c mice. |
| *C. freundii* | *Citrobacter freundii* P106E (preterm infant fecal isolate) |
| *E. cloacae* | *Enterobacter cloacae* ATCC13047 |
| S. *sonnei* | *Shigella sonnei* 53G-LVP |
| *S. enteritica* | *Salmonella enterica Enteriditis* ICC320 PT4 |
| *E. coli* Δ*ompA* | *E. coli* OmpA deletion mutant |
| *A. baumannii* | Acinetobacter baumannii ATCC17978 |

**Supplementary Table 2 conjugative plasmids**

| Plasmids | Description |
| --- | --- |
| R27 | Prototype plasmid for IncH |
| RA1 | Prototype plasmid for IncA/C |
| pABAY10001 | Plasmid expressing TraN_V_ isolated from *A. baumannii* |
| R27::RA1 tip | R27 expresses RA1 tip (AA 230C-360D) |
| RA1::R27 tip | RA1 expresses R27 tip (AA 305S-457E) |
| R27 Δtip | Tip-less mutant for R27 |
| RA1 Δtip | Tip-less mutant for RA1 |
| R27::TraN_L_γ | R27 expresses a TraN_L_γ cloned from a plasmid p-NDM-MAR (AA 304S-457D) |
| RA1::TraN_M_β | RA1 expresses a TraN_M_β cloned from a plasmid p-NDM-US (aa 205N-354V) |
| R27::TraN_V_ | R27 expresses the TraN_V_ (2 tips together) onto its TraN region (AA 280P-604K) which is cloned from pABAY10001 (AA 149V-628R) |
| R27::TraN_V_ (T1) | R27 expresses the TraN_V_ (T1)7 cloned from pABAY10001 (AA 158C-306T) |
| R27::TraN_TH_ (T2) | R27 expresses the TraN_V_ (T2)8 cloned from pABAY10001 (AA 371D-512Y) |

AA: amino acid

**Supplementary Table 3 Plasmids used for mutagenesis**

| Vector | Description |
| --- | --- |
| pACBSR | SmR; expresses I-SceI and lambda-red induced by L-Ara. |
| pSEVA612S | GmR; integrative plasmid (ori R6K) that harbours the oriT for tri-parental mating. |
| pSEVA612S_R27::RA1 tip | pSEVA612S derivative; swabs the RA1 tip onto R27 |
| pSEVA612S_RA1::R27 tip | pSEVA612S derivative; swabs the R27 tip onto RA1 |
| pSEVA612S_R27Δtip | pSEVA612S derivative; deletes the TraN tip region from R27 |
| pSEVA612S_RA1Δtip | pSEVA612S derivative; deletes the TraN tip region from RA1 |
| pSEVA612S_R27::TraN_Lγ_ tip | pSEVA612S derivative; substitutes the tip of TraN  in R27 with a chimeric TraN expressing tip of  TraN_Lγ_ |
| pSEVA612S_RA1::TraN_Mβ_ | pSEVA612S derivative; substitutes the tip of TraN  in RA1 with a chimeric TraN expressing tip of  TraN_Mβ_ |
| pSEVA612S_ R27::TraN_V_ | pSEVA612S derivative; substitutes the tip of TraN  in R27 with a chimeric TraN expressing tip of  TraN_V_ |
| pSEVA612S_R27::TraN_V_ (T1) | pSEVA612S derivative; substitutes the tip of TraN  in R27 with a chimeric TraN expressing tip of  TraN_V_ (tip 1) |
| pSEVA612S_R27::TraN_V_ (T2) | pSEVA612S derivative; substitutes the tip of TraN  in R27 with a chimeric TraN expressing tip of  TraN_V_ (tip 2) |

**Supplementary Table 4 Primers used in this study**

| Primers | Sequences (5’-3’) | Description |
| --- | --- | --- |
| pSEVA612S_F | ATTACCCTGTTATCCCTATACTG | Amplifies linear pSEVA612S |
| pSEVA612S_R | TAGGGATAACAGGGTAATCCG |  |
| UP R27 HR_fwd | gttatccctaCCGGGGAAAATCTGAGCG | Amplifies the upstream homology region of R27 TraN |
| UP R27 HR_rev | ttattcccaaAAAATCACGAGTTATCGTGCAAC |  |
| RA1 TIP_fwd | tcgtgattttTTGGGAATAACAAGTGATGTTC | Amplifies the TraN tip region from RA1 |
| RA1 TIP_rev | tatgtttatcGTCCTCAACTTGCTTGGTG |  |
| R27downHR_fwd | agttgaggacGATAAACATATACAGGAGCCCGC | Amplifies the downstream homology region of R27 TraN |
| R27 downHR_rev | gttatccctaCGCGGCCACTTTCCCAAAATC |  |
| RA1 UPHR_fwd | gttatccctaCTATTTCAATGCCTACGTTG | Amplifies the upstream homology region of RA1 TraN |
| RA1 UPHR_rev | ccggcacggaGGAATGGATAATATCGCAATC |  |
| R27 TIP_fwd | tatccattccTCCGTGCCGGTTTATATC | Amplifies the TraN tip region from R27 |
| R27 TIP_rev | gagtccattgCTCCAACTTCATATTTTCAAAAG |  |
| RA1downHR_fwd | gaagttggagCAATGGACTCCGCAGACTTG | Amplifies the downstream homology region of RA1 TraN |
| RA1 downHR_rev | gttatccctaTCTGGCGAAATCGGTACTC |  |
| ndm mar tip_fwd | tcgtgattttagtgtaccagtttacatctc | Amplifies the TraN tip region from TraN_Lγ_ |
| ndm mar tip_rev | tatgtttatcatcttcaagtttgatattttcaaaaattag |  |
| NDM-US tip_fwd | tatccattcctctgtggtgaagcactatg | Amplifies the TraN tip region from TraN_Mβ_ |
| NDM-US tip_rev | gagtccattgggctttcgatgggtcatag |  |
| RA1 TraN tip ext-f | ccgcgttgccggtacatggtta | Check the substitution of R27 TraN tip/TraN_M_β |
| RA1 TraN tip ext-r | taccgcgccatctagcttggga |  |
| RA1 TraN tip san-f | cgcaacaaatcgttctcgcccg | For Sanger Sequencing |
| RA1 TraN tip san-r | ccagggcaaatgctggagccat |  |
| R27 TraN tip ext-f | cggccagcattatcgcctccag | Check the substitution of RA1 TraN tip/TraN_V_/TraN_L_β |
| R27 TraN tip ext-r | ccaatcccaacgccaaccggaa |  |
| R27 TraN tip san-f | gcacactttccgggagtcgctt | For Sanger Sequencing |
| R27 TraN tip san-r | cagttcaatgctggtggccggt |  |
| abay10001_fwd | tacgcactatgtcaaacctgtctatgaag | Amplifies the 2 tips region from TraN_V_ |
| abay10001_rev | tgactacgtcacgtgctggtgttttttc |  |
| tip 1_fwd | tatgtttatctgtatgcaggggcaagctttc | Amplifies the tip 1 region from TraN_V_ |
| tip 1_rev | tcgtgattttggtattaaatacaatcaactcaagtgtattttcac |  |
| tip 2_fwd | tatgtttatcgacagcaacttctatcaac | Amplifies the tip 2 region from TraN_V_ |
| tip 2_rev | tcgtgattttatacaatctgattgtatttgatcc |  |
